# Reconstructing contact network parameters from viral phylogenies

**DOI:** 10.1101/050435

**Authors:** Rosemary M. McCloskey, Richard H. Liang, Art F.Y. Poon

## Abstract

Models of the spread of disease in a population often make the simplifying assumption that the population is homogeneously mixed, or is divided into homogeneously mixed compartments. However, human populations have complex structures formed by social contacts, which can have a significant influence on the rate of epidemic spread. Contact network models capture this structure by explicitly representing each contact which could possibly lead to a transmission. We developed a method based on kernel approximate Bayesian computation (kernel-ABC) for estimating structural parameters of the contact network underlying an observed viral phylogeny. The method combines adaptive sequential Monte Carlo for ABC, Gillespie simulation for propagating epidemics though networks, and a kernel-based tree similarity score. We used the method to fit the Barabási-Albert network model to simulated transmission trees, and also applied it to viral phylogenies estimated from five published HIV sequence datasets. On simulated data, we found that the preferential attachment power and the number of infected nodes in the network can often be accurately estimated. On the other hand, the mean degree of the network, as well as the total number of nodes, were not estimable with kernel-ABC. We observed substantial heterogeneity in the parameter estimates on real datasets, with point estimates for the preferential attachment power ranging from 0.06 to 1.05. These results underscore the importance of considering contact structures when performing phylodynamic inference. Our method offers the potential to quantitatively investigate the contact network structure underlying viral epidemics.

## Introduction

When an infectious disease spreads through a population, transmissions are generally more likely to occur between certain pairs of individuals. Such pairs must have a particular mode of contact with one another, which varies with the mode of transmission of the disease. For airborne pathogens, physical proximity may be sufficient, while for sexually transmitted diseases, sexual or in some cases blood-to-blood contact is required. The population together with the set of links between individuals along which transmission can occur is called the contact network [26, 34]. The structure of the contact network underlying an epidemic can profoundly impact the speed and pattern of the epidemic’s expansion. Network structure can influence the prevalence trajectory [39, 33] and epidemic threshold [2], in turn affecting the estimates of quantities such as effective population size [18]. From a public health perspective, contact networks have been explored as tools for curtailing epidemic spread, by way of interventions targeted to well-connected nodes [59]. True contact networks are a challenging type of data to collect, requiring extensive epidemiological investigation [60].

Viral sequence data, on the other hand, has become relatively inexpensive and straightforward to collect on a population level. Due to the high mutation rate of RNA viruses, epidemiological processes impact the course of viral evolution, thereby shaping the inter-host viral phylogeny [14]. The term “phylodynamics” was coined to describe this interaction, as well as the growing family of inference methods to estimate epidemiological parameters from viral phylogenies [20]. These methods have revealed diverse properties of local viral outbreaks, from basic reproductive number [51], to the degree of clustering [23], to the elevated transmission risk during acute infection [57]. On the other hand, although sophisticated methods have been developed for fitting complex population genetic models to phylogenies [56, 45], inference of structural network parameters has to date been limited. However, it has been shown that network structure has a tangible impact on phylogeny shape [28, 9, 18, 46, 54], suggesting that such statistical inference might be possible [60].

Survey-based studies of sexual networks [8, 30, 47] have found that these networks tend to have a degree distribution which follows a power law [although there has been some disagreement, see 22]. That is, the number of nodes of degree *k* is proportional to *k*^−*γ*^ for some constant *γ*. These networks are also referred to as “scale-free” [1]. One process by which scale-free networks can be generated is preferential attachment, where nodes with a high number of contacts attract new connections at an elevated rate. The first contact network model incorporating preferential attachment was introduced by Barabási and Albert [1], and is now referred to as the Barabási-Albert (BA) model. Under this model, networks are formed by iteratively adding nodes with *m* new edges each. In the most commonly studied formulation, these new edges are joined to existing nodes of degree *k* with probability proportional to *k*, so that nodes of high degree tend to attract more connections. Barabási and Albert suggested an extension where the probability of attaching to a node of degree *k* is *k*^*α*^ for some non-negative constant *α*, and we use this extension in this work.

Previous work offers precedent for the possibility of statistical inference of structural network parameters. Britton and O’Neill [5] develop a Bayesian approach to estimate the edge density in an Erdős-Rényi network [16] given observed infection dates, and optionally recovery dates. Their approach was later extended by Groendyke, Welch, and Hunter [21] and applied to a much larger data set of 188 individuals. Volz and Meyers [58] and Volz [55] developed differential equations describing the spread of a susceptible-infected (SI) epidemic on static and dynamic contact networks with several degree distributions, which could in principle be used for inference if observed incidence trajectories were available. Brown et al. [6] analysed the degree distribution of an approximate transmission network, estimated based on genetic similarity and estimated times of infection, relating 60% of HIV-infected men who have sex with men (MSM) in the United Kingdom. The transmission network is a subgraph of the contact network which includes only those edges which have already led to a new infection. The authors found that a Waring distribution, which is produced by a more sophisticated preferential attachment model, was a good fit to their estimated network.

Standard methods of model fitting involve calculation of the likelihood of observed data under the model. In maximum likelihood estimation, a quantity proportional to the likelihood is optimized, often through a standard multi-dimensional numerical optimization procedure. Bayesian methods integrate prior information by optimizing the posterior probability instead. To avoid calculation of a normalizing constant, Bayesian inference is often performed using Markov chain Monte Carlo (MCMC), which uses likelihood *ratios* in which the normalizing constants cancel out. Unfortunately, it is generally difficult to explicitly calculate the likelihood of an observed transmission tree under a contact network model, even up to a normalizing constant. To do so, it would be necessary to integrate over all possible networks, and also over all possible labellings of the internal nodes of the transmission tree. While it is not known (to us) whether such integration is tractable, a simpler alternative is offered by likelihood-free methods, namely approximate Bayesian computation (ABC) [53, 3]. ABC leverages the fact that, although calculating the likelihood may be impractical, generating simulated datasets according to a model is often straightforward. If our model fits the data well, the simulated data it produces should be similar to the observed data. More formally, if *D* is the observed data, the posterior distribution *f*(*θ* | *D*) on model parameters *θ* is replaced as the target of statistical inference by *f*(*θ* | *ρ*(*D*̂,*D*) < *ε*), where *ρ* is a distance function, *D*̂ is a simulated dataset according to *θ*, and *ε* is a small tolerance [52]. In the specific case when *ρ* is a kernel function, the approach is known as kernel-ABC [35, 41].

Here, we develop a method using kernel-ABC to estimate the parameters of contact network models from observed phylogenetic data. The distance function we use is the tree kernel developed by Poon et al. [42], which computes a weighted dot product of the trees’ representations in the space of all possible subset trees. We apply the method to investigate the parameters of the BA network model on a variety of simulated and real datasets. Our results show that some network parameters can be inferred with reasonable accuracy, while others have a minimal detectable impact on tree shape and therefore cannot be estimated accurately. We also find that these parameters can vary considerably between real epidemics from different settings.

## Methods

### *Netabc*: phylogenetic inference of contact network parameters with kernel-ABC

We have developed a kernel-ABC-based method to perform statistical inference of contact network parameters from a transmission tree estimated from an observed viral phylogeny. We implemented the adaptive sequential Monte Carlo (SMC) algorithm for ABC developed by Del Moral, Doucet, and Jasra [12]. The SMC algorithm keeps track of a population of parameter “particles”, which are initially sampled from the parameters’ joint prior distribution. Several datasets are simulated under the model of interest for each of the particles. In this case, the datasets are transmission trees, which are generated by a two-step process. First, a contact network is simulated according to the network model being fit. Second, a transmission tree is simulated over that network with a Gillespie simulation algorithm [17], in the same fashion as several previous studies [*e.g.* 46, 28]. The particles are weighted according to the similarity between their associated simulated trees and the observed tree. To quantify this similarity, we used the tree kernel developed by Poon et al. [42]. Particles are iteratively perturbed by applying a Metropolis-Hastings kernel and, if the move is accepted, simulating new datasets under the new parameters. When a particle’s weight drops to zero, because its simulated trees are too dissimilar to the observed tree, the particle is dropped from the population, and eventually replaced by a resampled particle with a higher weight. As the algorithm progresses, the population converges to a Monte Carlo approximation of the ABC target distribution, which is assumed to approximate the desired posterior [12, 52]. A computer program implementing our method is freely available at https://github.com/rmcclosk/netabc (last accessed April 26, 2016).

To check that our implementation of Gillespie simulation was correct, we reproduced Figure 1A of Leventhal et al. [28] (our fig. S1), which plots the unbalancedness of transmission trees simulated over four network models at various levels of pathogen transmissibility. Our implementation of adaptive ABC-SMC was tested by applying it to the same mixture of Gaussians used by Del Moral, Doucet, and Jasra to demonstrate their method (originally used by Sisson, Fan, and Tanaka [49]). We were able to obtain a close approximation to the function (see fig. S2), and attained the stopping condition used by the authors in a comparable number of steps.

**Figure 1.**
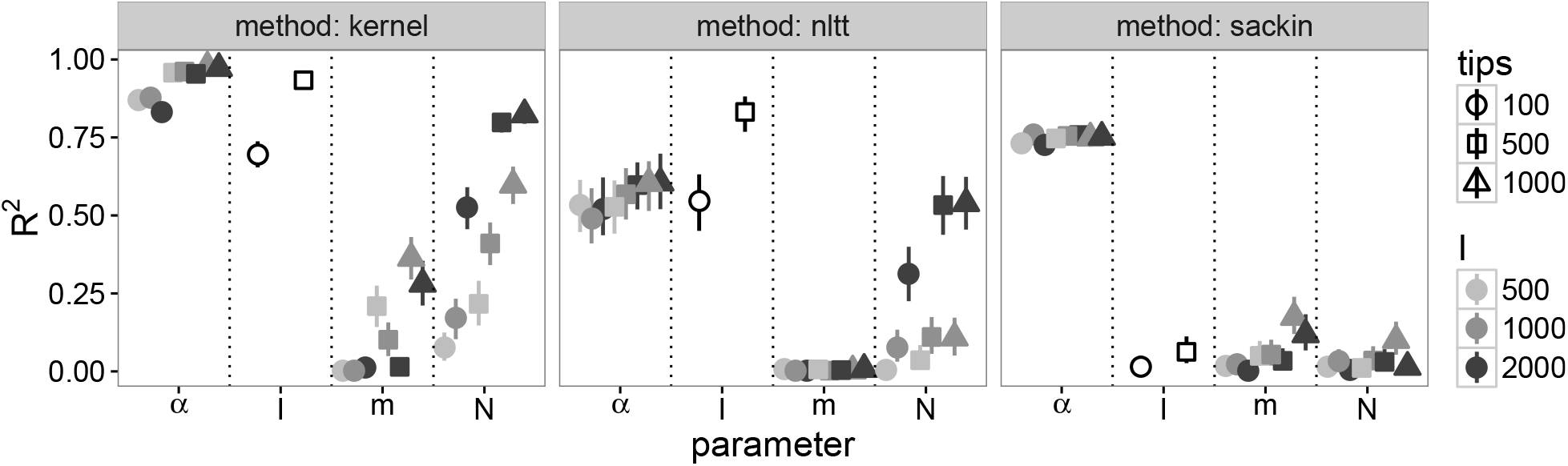
Cross-validation accuracy of kernel-SVR classifier (left), SVR classifier using nLTT (centre), and linear regression using Sackin’s index (right) for BA model parameters. Kernel meta-parameters were set to *γ* = 0.3 and *σ* = 4. Each point was calculated based on 300 simulated transmission trees over networks with three different values of the parameter being tested. Vertical lines are empirical 95% confidence intervals based on 1000 two-fold cross-validations.

Nodes in our networks followed simple SI dynamics, meaning that they became infected at a rate proportional to their number of infected neighbours, and never recovered. For all analyses, the transmission trees’ branch lengths were scaled by dividing by their mean. We used the *igraph* library’s implementation of the BA model [10] to generate the graphs. The analyses were run on Westgrid (https://www.westgrid.ca/) and a local computer cluster.

### Kernel classifiers

We used the phylogenetic kernel developed by Poon et al. [42] to test whether the parameters of the BA model had an effect on tree shape. 100 networks were simulated under each of three different values of *α*: 0.5, 1.0, and 1.5 (300 networks total). The other parameters were fixed to the following values: *N* = 5000, *I* = 1000, and *m* = 2. A transmission tree with 500 tips was simulated over each network (300 transmission trees total). The 300 trees were compared pairwise with the tree kernel to form a 300 × 300 kernel matrix. The kernel meta-parameters *λ* (the “decay factor”), and *σ* (the “radial basis function variance”) [see 42], were set to 0.3 and 4 respectively. We constructed a kSVR classifier for *α* using the *kernlab* package [62], and evaluated its accuracy with 1000 two-fold cross-validations.

Three similar experiments were performed for the other BA model parameters (one experiment per parameter). *m* was varied between 2, 3, and 4; *I* between 500, 1000, and 2000; and *N* between 3000, 5000, and 8000. The parameters not being tested were fixed at the values *N* = 5000, *I* = 1000, *m* = 2, and *α* = 1. Thus, we performed a total of four kSVR cross-validations, one for each of the BA model parameters *α*, *I*, *m*, and *N*. We repeated these four cross-validations with different values of *λ* (0.2,0.3, and 0.4) and *σ* (2^−3^, 2^−2^, 2^3^), as well as on trees with differing numbers of tips (100, 500, and 1000) and in epidemics of differing size (500, 1000, and 2000). The combination of the number of sampled individuals (*i.e*. the number of tips) and the epidemic size (*i.e. I*) will be referred to as an “epidemic scenario”. When evaluating the classifier for *I*, we did not consider trees with 1000 tips, because one of the tested *I* values was 500, and the number of tips cannot be larger than *I*.

For each of the four parameters, we also tested a linear regression against Sackin’s index [48] and an ordinary SVR based on the normalized lineages-through-time (nLTT) statistic [24].

### ABC simulations

We simulated three transmission trees, each with 500 tips, under every element of the Cartesian product of these parameter values: *N* = 5000, *I* = {1000, 2000}, *m* = {2, 3, 4}, and *α* = {0.0, 0.5, 1, 1.5}. This produced a total of 24 parameter combinations × three trees per combination = 72 trees total. The adaptive ABC algorithm was applied to each tree with these priors: *m* ~ DiscreteUniform(1, 5), *α* ~ Uniform(0, 2), and (*N*, *I*) jointly uniform on the region {500 ≤ *N* ≤ 15000, 500 ≤ *I* ≤ 5000, *I* ≤ *N*}. Proposals for *α*, *N*, and *I* were Gaussian, while proposals for *m* were Poisson. Following Del Moral, Doucet, and Jasra [12] and Beaumont et al. [4], the variance of all proposals was equal to the empirical variance of the particles.

The algorithm was run with 1000 particles, 5 simulated datasets per particle, and the “quality” parameter controlling the decay rate of the tolerance *ε* set to 0.95. We used the same stopping criterion as Del Moral, Doucet, and Jasra, namely when the MCMC acceptance rate dropped below 1.5%. Point estimates for the parameters were obtained by taking the highest point of an estimated kernel density on the final set of particles, calculated using the *density* function with the default parameters in *R*. Highest posterior density (HPD) intervals were calculated with the *HPDinterval* function from the *R* package *coda* [40].

Two further analyses were performed to address potential sources of error. To evaluate the effect of model misspecification in the case of heterogeneity among nodes, we generated a network where half the nodes were attached with power *α* = 0.5, and the other half with power *α* = 1.5. The other parameters for this network were *N* = 5000, *I* = 1000, and *m* = 2. To investigate the effects of potential sampling bias, we simulated a transmission tree where the tips were sampled in a peer-driven fashion, rather than at random. That is, the probability to sample a node was twice as high if any of that node’s network peers had already been sampled. The parameters of this network were *N* = 5000, *I* = 2000, *m* = 2, and *α* = 0.5.

### Investigation of published data

We applied our kernel-ABC method to several published HIV datasets. Because the BA model generates networks with a single connected component, we specifically searched for datasets which originated from existing clusters, either phylogenetically or geographically defined. Characteristics of the datasets we investigated are given in table 1.

**Table 1.**
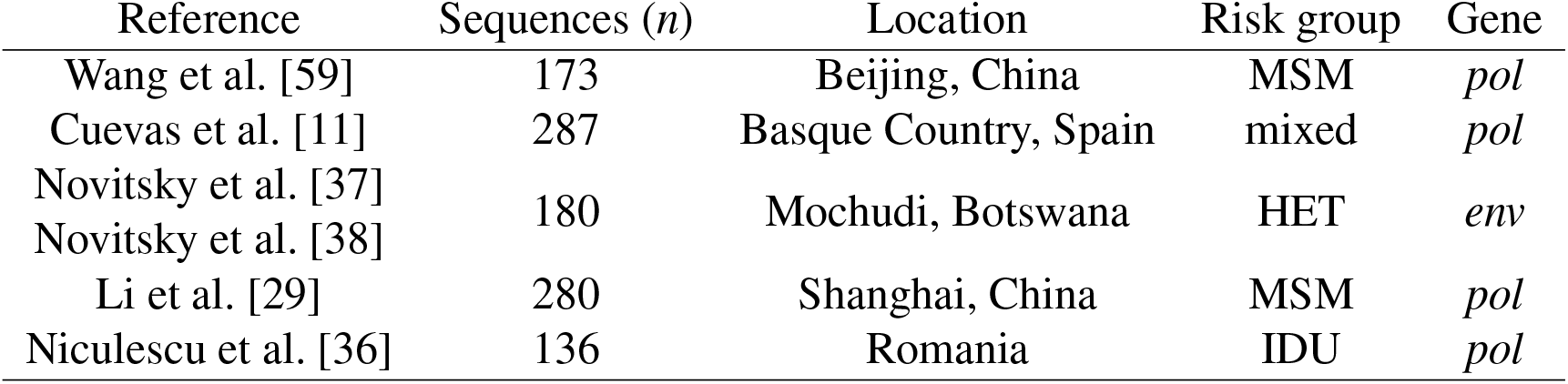
Characteristics of published datasets investigated with kernel-ABC. Acronyms: MSM, men who have sex with men; IDU, injection drug users; HET, heterosexual. The Novitsky et al. [37] and Novitsky et al. [38] data was sampled from a primarily heterosexual risk environment, but did not explicitly exclude other risk groups.

We downloaded all sequences associated with each study from GenBank. For the Novitsky et al. [38] data, each sequence was aligned pairwise to the HXB2 reference sequence (Genbank accession number K03455) and the hypervariable regions were clipped out with *BioPython* version 1.66+ [7]. Sequences were multiply aligned using *MUSCLE* version 3.8.31 [15], and alignments were manually inspected with *Seaview* version 4.4.2 [19]. Phylogenies were constructed from the nucleotide alignments by approximate maximum likelihood using *FastTree2* version 2.1.7 [44] with the generalized time-reversible model. Transmission trees were estimated by rooting and time-scaling the phylogenies by root-to-tip regression, using a modified version of Path-O-Gen (distributed as part of BEAST [13]) as described previously [41].

Two of the datasets [29, 38] were initially much larger than the others, containing 1265 and 1299 sequences respectively. To ensure that the analyses were comparable, we reduced these to a number of sequences similar to the smaller datasets. For the Li et al. [29] data, we detected a cluster of size 280 using a patristic distance cutoff of 0.02 as described previously [43]. Only sequences within this cluster were carried forward. For the Novitsky et al. [38] data, no large clusters were detected using the same cutoff, so we analysed a subtree of size 180 chosen arbitrarily.

For all datasets, we used the priors *α* ~ Uniform(0, 2) and *N* and *I* jointly uniform on the region {*n* ≤ *N* ≤ 10000, *n* ≤ *I* ≤ 10000, *I* ≤ *N*}, where *n* is the number of tips in the tree. Since the value *m* = 1 produces networks with no cycles, which we considered fairly implausible, we ran one analysis with the prior *m* ~ DiscreteUniform(1, 5), and one with the prior *m* ~ DiscreteUniform(2, 5). The other parameters to the SMC algorithm were the same as used for the simulation experiments.

## Results

### Kernel classifiers

We investigated the parameters of the BA network model [1]. In addition to *m* and *α* (see Introduction), we considered *N*, which denotes the total number of nodes in the network, and *I*, which is the number of infected nodes at which to stop the simulation and sample the transmission tree. To examine the effect of these parameters on tree shape, we simulated transmission trees under different parameter values, calculated pairwise tree kernel scores between them, and attempted to classify the trees using a kernel support vector machine (kSVR). We also tested classifiers based on Sackin’s index [48] and the normalized lineages-through-time (nLTT) statistic [24]. Accuracy of the kSVRs varied based on the parameter being tested (fig. 1, left). Classifiers based on two other tree statistics, the nLTT and Sackin’s index, generally exhibited worse performance than the tree kernel, although the magnitude of the disparity varied between the parameters (fig. 1, centre and right). The results were largely robust to variations in the tree kernel meta-parameters *λ* and *σ* (figs. S3 to S6).

When classifying *α*, the kernel-SVR classifier had an average *R*^2^ of 0.92, compared to 0.56 for the nLTT-based SVR, and 0.75 for the linear regression against Sackin’s index. There was little variation about the mean for different tree and epidemic sizes. No classifier could accurately identify the *m* parameter in any epidemic scenario, with average *R*^2^ values of 0.12 for kSVR, 0.01 for the nLTT, and 0.06 for Sackin’s index. Again, there was little variation in accuracy between epidemic scenarios, although the accuracy of the kSVR was slightly higher on 1000-tip trees (fig. 1, left).

The accuracy of classifiers for *I* varied significantly with the number of tips in the tree. For 100-tip trees, the average *R*^2^ values were 0.7, 0.55, and 0.02 for the tree kernel, nLTT, and Sackin’s index respectively. For 500-tip trees, the values increased to 0.93, 0.83, and 0.07. Finally, the performance of classifiers for *N* depended heavily on the epidemic scenario. The *R*^2^ of the kSVR classifier ranged from 0.08 for the smallest epidemic and smallest sample size, to 0.82 for the largest. Likewise, *R*^2^ for the nLTT-based SVR ranged from 0.01 to 0.54. Sackin’s index did not accurately classify *N* in any scenario, with an average *R*^2^ of 0.03 and little variation between scenarios.

### ABC simulations

Figure 2 shows maximum *a posteriori* (MAP) point estimates of the BA model parameters obtained with kernel-ABC on simulated data. The estimates shown correspond only to the simulations where the *m* parameter was set to 2, however the results for *m* = 3 and *m* = 4 were similar (figs. S7 and S8). Average boundaries of 95% HPD intervals are given in table 2.

**Figure 2.**
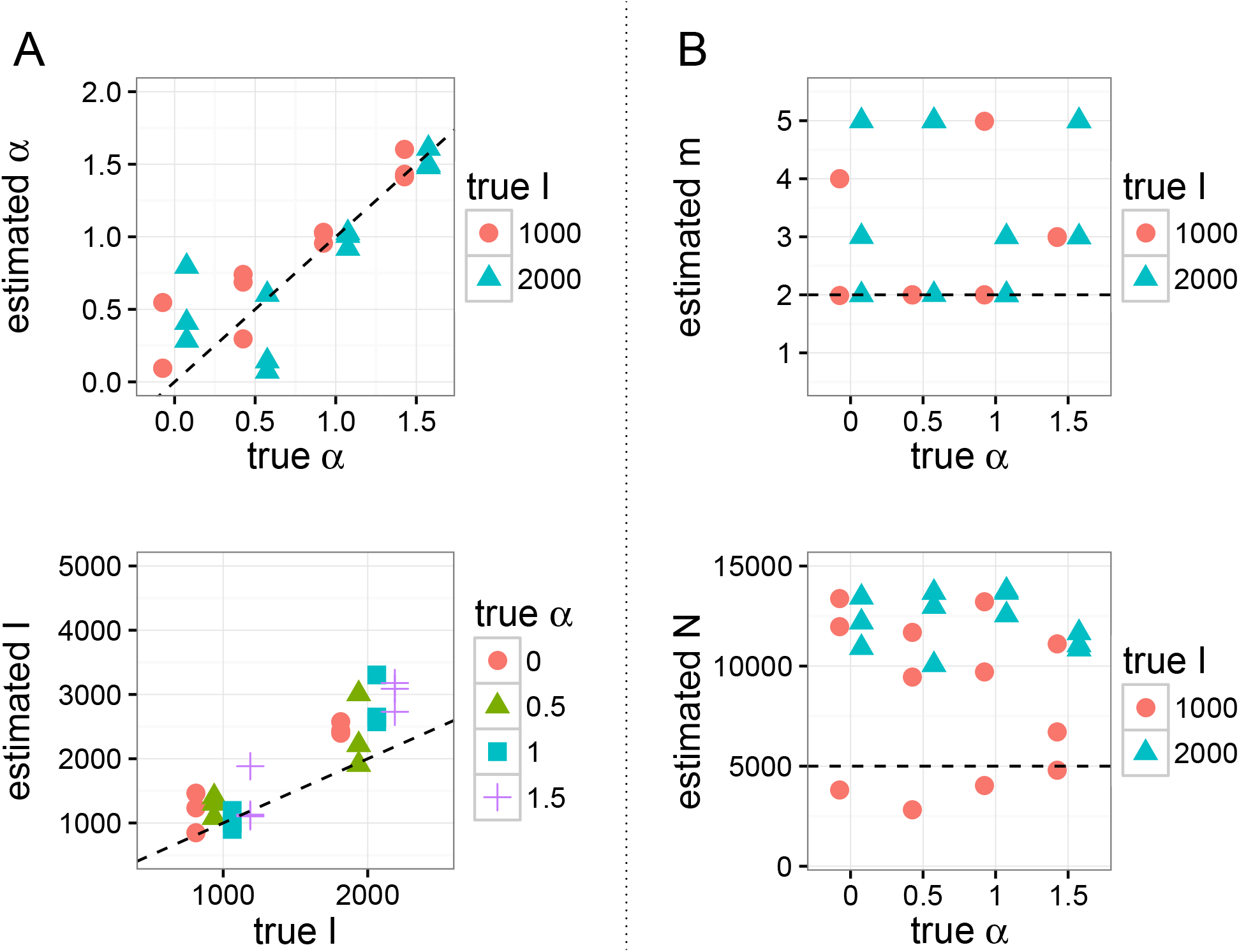
Point estimates of BA model parameters obtained by running kernel-ABC on simulated phylogenies without training, for simulations with *m* = 2. Dotted lines indicate true values, and limits of the *y*-axes are regions of uniform prior density. (A) Estimates for *α* and *I* against their true values in simulations. (B) Estimates for *m* and *N*, which were held fixed in these simulations, against true values of *α*.

**Table 2.**
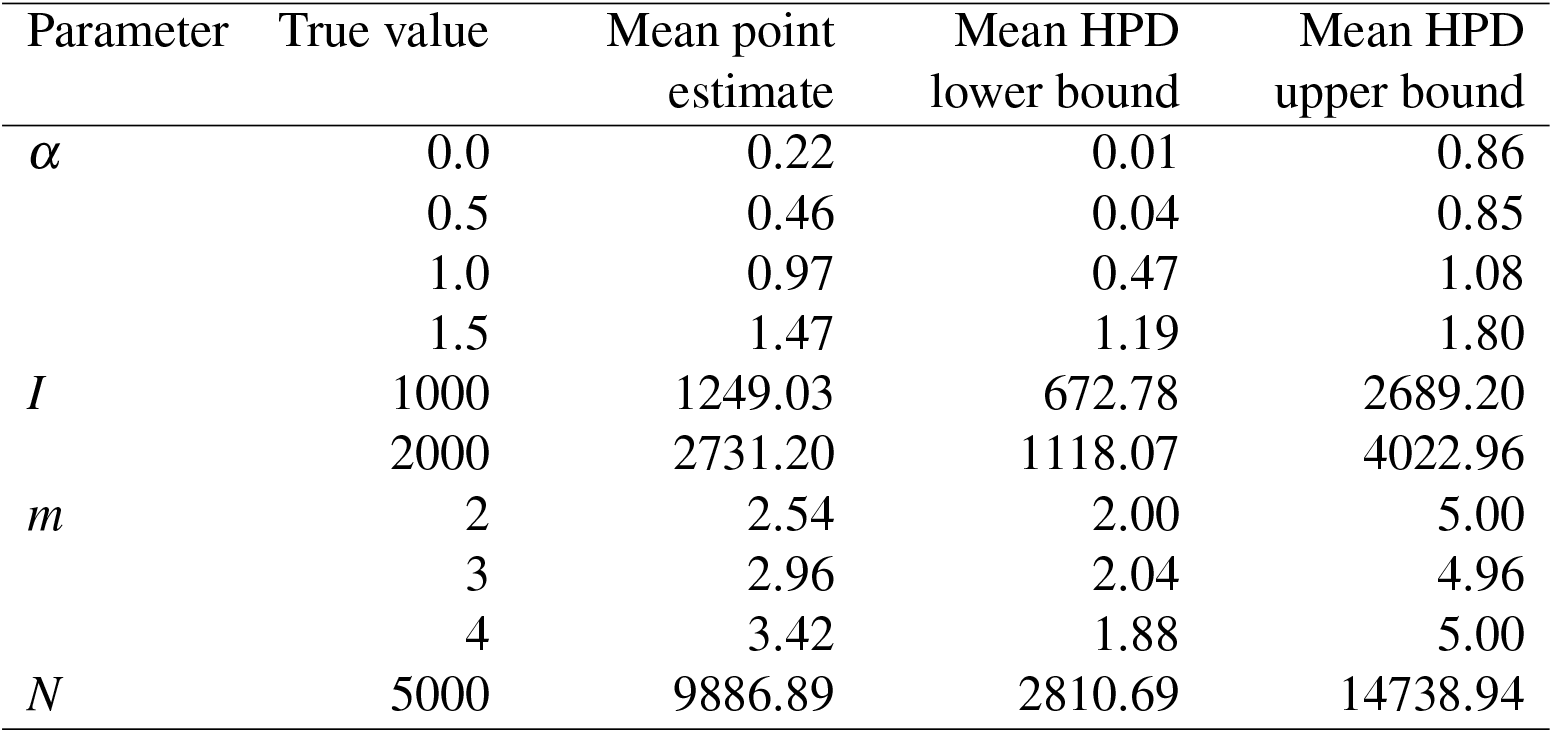
Average maximum *a posteriori* point estimates and 95% highest posterior density (HPD) interval widths for BA model parameter estimates obtained with kernel-ABC. Three transmission trees were simulated under each combination of the listed parameter values, and the parameters were estimated with kernel-ABC without training.

The accuracy of the parameter estimates obtained with kernel-ABC paralleled the results from the kSVR classifier. Of the four parameters, *α* was the most accurately estimated, with point estimates having a median [IQR] absolute error of 0.08 [0.05 - 0.17]. The errors when the true value of *α* was zero were significantly greater than those for the other values (Wilcoxon rank-sum test, *p* = 0.0078). Errors in estimating *α* also varied with the true value of *m* just at the threshold of statistical significance (*p* = 0.05), but did not vary across the true values of *N* or *I* (both one-way ANOVA). Estimates for *I* were relatively accurate, with point estimate errors of 395 [207 - 683] individuals. These errors were significantly higher when the true value of *α* was at least 1 (Wilcoxon rank-sum test, *p* = 0.0077) and when the true value of *I* was 2000 (*p* < 10^−5^). The true value of *m* did not affect the estimates of *I* (one-way ANOVA).

The *m* parameter was estimated correctly in only 27 % of simulations, barely better than random guessing. The true values of the other parameters did not significantly affect the estimates of *m* (both one-way ANOVA). Finally, the total number of nodes *N* was consistently over-estimated by about a factor of two (error 5987 [2060 - 7999] individuals). No parameters influenced the accuracy of the *N* estimates (all one-way ANOVA).

The dispersion of the ABC approximation to the posterior also varied between the parameters, with narrower HPD intervals for the parameters with more accurate point estimates (table 2). Figure 3 shows the distributions for for one simulation. Equivalent plots for one replicate simulation with each studied parameter combination can be found in figs. S18 to S41. HPD intervals around *α* and *I* were often narrow relative to the region of nonzero prior density, whereas the intervals for *m* and *N* were more widely dispersed.

**Figure 3.**
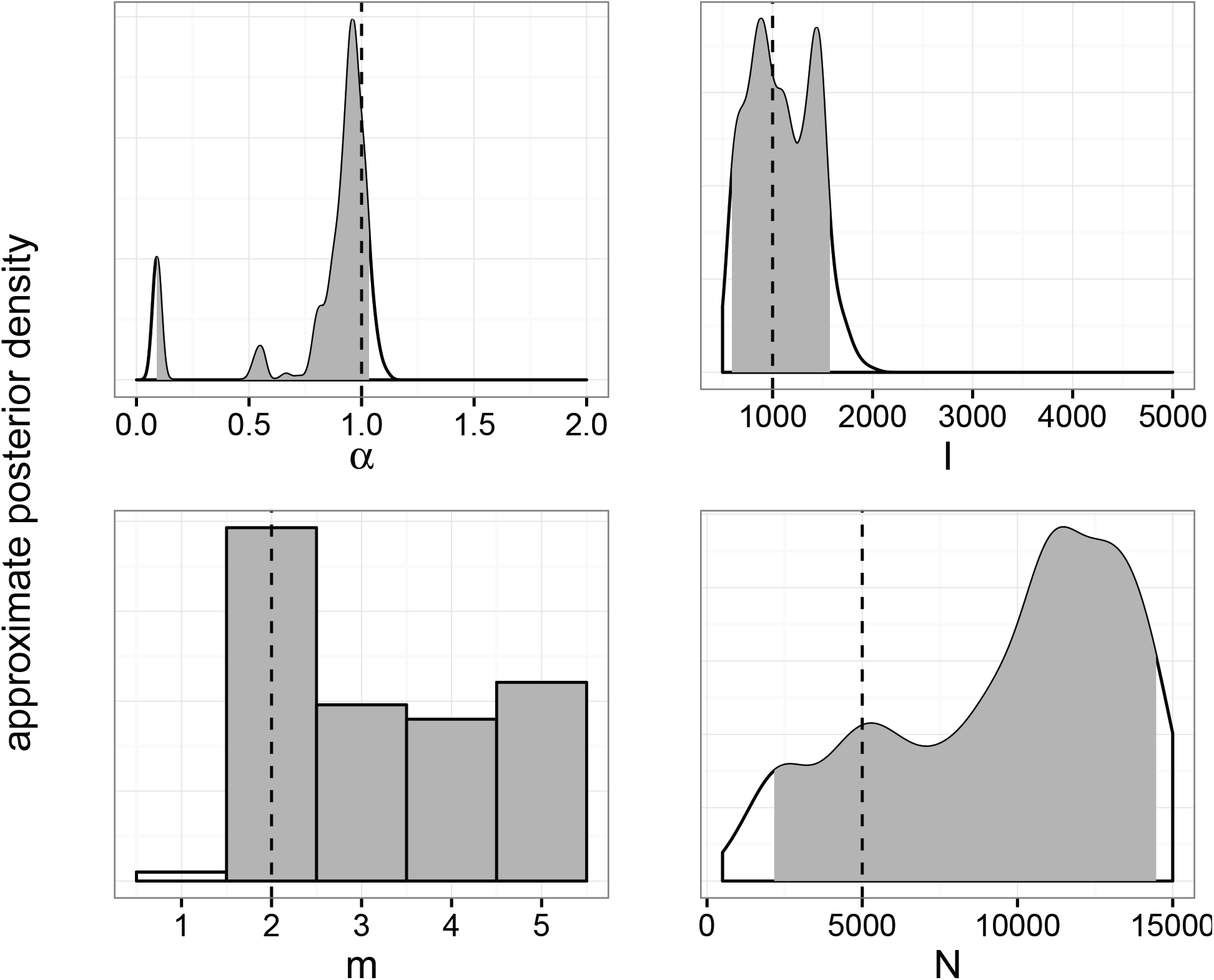
Marginal posterior distributions of BA model parameters estimated with kernel-ABC for a single simulated transmission tree. Dotted lines and shaded polygon indicate true values.

To test the effect of model misspecification, we simulated one network where the nodes exhibited heterogeneous preferential attachment power (half 0.5, the other half 1.5), with *m* = *2*, *N* = 5000, and *I* = 1000. The MAP [95% HPD] estimates for each parameter were: *α*, 1.1 [0.6 - 1.16]; *I*, 1116 [662 - 4455]; *m*, 3 [1 - 5]; *N*, 12657 [3590 - 14977]. The approximate posterior distributions for this simulation are shown in fig. S10. To test the effect of sampling bias, we sampled one transmission tree in a peer-driven fashion, where the probability to sample a node was twice as high if one of its peers had already been sampled. The parameters for this experiment were *N* = 5000, *m* = 2, *α* = 0.5, and *I* = 2000. The estimated values were *α*, 0.23 [0.02 - 0.64]; *I*, 2407 [1419 - 3838]; *m*, 3 [2 - 5]; *N*, 8716 [2741 - 14615]. The approximate posterior distributions are shown in fig. S11. Both of these results were in line with estimates obtained on other simulated datasets (table 2), although the estimate of peer-driven sampling for *α* was somewhat lower than typical.

### Real data

We applied kernel-ABC to five published HIV datasets (table 1), and found substantial heterogeneity among the parameter estimates (figs. 4 and S12). Plots of the marginal posterior distributions for each dataset are shown in figs. S13 to S17. Two of the datasets [36, 59] had estimated *α* values near unity for the prior allowing *m* = 1 (MAP estimates [95% HPD] 1.05 [0.04 - 1.27] and 0.84 [0.01 - 1.02] respectively). The MAP estimates did not change appreciably when *m* = 1 was disallowed by the prior, although the credible interval of the Niculescu et al. [36] data was narrower (0.04 - 1.27). When *m* = 1 was permitted, the Li et al. [29] and Cuevas et al. [11] both had low estimated *α* values (0.06 [0.01 - 0.73] and 0.19 [0.01 - 0.8]). However, the MAP estimates increased when *m* = 1 was not permitted, although the HPD intervals remained roughly the same (0.78 [0.02 - 0.94] and 0.59 [0.07 - 0.95]). The Novitsky et al. [38] data had a fairly low estimated *α* for both priors on *m* (0.32 for *m* ≥ 1; 0.39 for *m* ≥ 2). However, the confidence interval was much wider when *m* = 1 was allowed ([0.04 −1.62] for *m* ≥ 1 vs. 0 - 0.73 for *m* ≥ 2).

**Figure 4.**
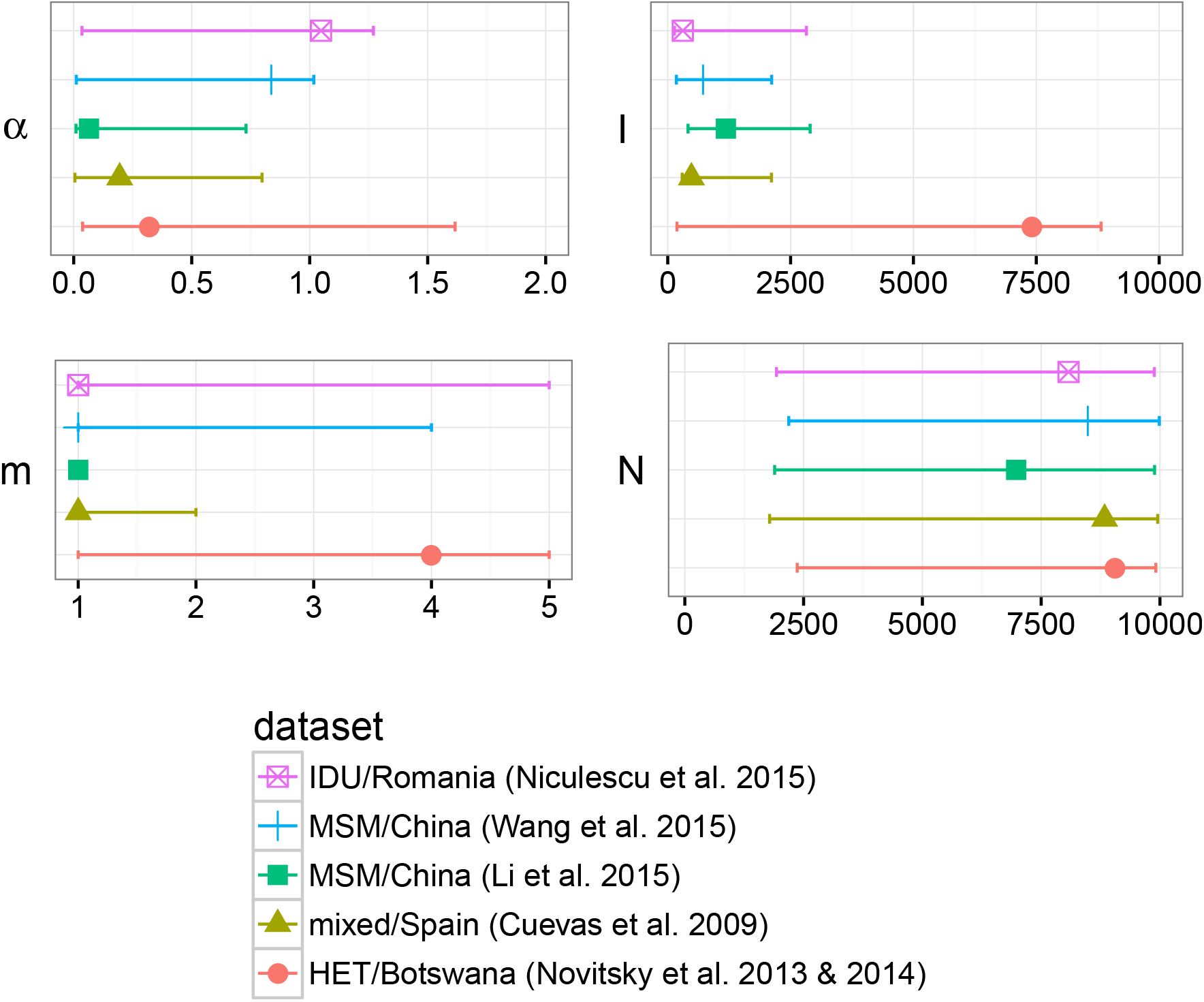
Maximum *a posteriori* point estimates and 95% HPD intervals for parameters of the BA network model, fitted to five published HIV datasets with kernel-ABC. *x*-axes indicate regions of nonzero prior density. In particular, the prior on *m* was DiscreteUniform(1, 5).

For all the datasets except Novitsky et al., estimated values of *I* were below 2000 when *m* = 1 was allowed, with relatively narrow HPD intervals compared to the nonzero prior density region (Cuevas et al., 482 [293 - 2111]; Niculescu et al., 307 [136 - 2822]; Li et al., 1183 [413 - 2897]; Wang et al., 719 [176 - 2114]). The Novitsky et al. data was the outlier, with a very high estimated *I*, and HPD interval spanning almost the entire prior region (7409 [187 - 8819]). The *I* estimates and HPD intervals were generally robust to the choice of prior on *m*, with slightly narrower HPD intervals (compare figs. 4 and S12).

The MAP estimate of *m* was equal to 1 for all but the Novitsky et al. data, when this value was allowed. However, the upper bound of the HPD interval was different for each dataset (Niculescu et al., 5; Wang et al., 4; Li et al., 1; Cuevas et al., 2). When *m* = 1 was disallowed, the MAP for all datasets was either 2 or 3, with HPD intervals spanning the entire prior region. The estimates for the total number of nodes *N* were largely uninformative for all samples, with almost all MAP estimates greater than 7500 and HPD intervals spanning almost the entire nonzero prior density region. The only exception was the Li et al. data, for which the MAP estimate was lower (6973) when *m* = 1 was allowed.

## Discussion

Contact networks can have a strong influence on epidemic progression, and are potentially useful as a public health tool [59, 32]. Despite this, few methods exist for investigating contact network parameters in a phylodynamic framework [although see 21, 55, 6, 28, for related work]. Kernel-ABC is a model-agnostic method which can be used to investigate any quantity that affects tree shape [41]. In this work, we developed a kernel-ABC-based method to infer the parameters of a contact network model. The method is general, and could be applied to any model from which contact networks can be simulated. We demonstrated the method on the BA model, which is a simple preferential attachment model giving rise to the power law degree distributions commonly observed in real-world networks.

By training a kernel-SVR classifier, we found that the *α* and *I* parameters, representing preferential attachment power and number of infected nodes, had a strong influence on tree shape. This was reflected in the relative accuracy of the kernel-ABC estimates of these parameters. The total number of nodes *N* had a weak influence on tree shape, which was most prominent when the epidemic size *I* and number of sampled tips were both large. The *m* parameter, representing the number of edges created in the network per vertex, did not produce much variation in tree shape, resulting in in both poorly performing classifiers and uninformative kernel-ABC estimates.

*N* was almost always significantly over-estimated using kernel-ABC. Since the prior on *N* and *I* is jointly uniform on a non-rectangular region (*I* ≤ *N*), there is more prior mass on high *N* values. In retrospect, it is unreasonable to expect good estimation of *N*, because adding more nodes to a BA network does not change the edge density or overall shape. This can be illustrated by imagining that we add a small number of nodes to a network after the epidemic simulation has already been completed. It is possible that none of these new nodes attains a connection to any infected node. Thus, running the simulation again on the new, larger network could produce the exact same transmission tree as before. We note also that our accurate estimates of *I* may have been influenced by this prior, which places more mass on low *I* values. However, the MAP estimate of *I* was very high for the Novitsky et al. [37] and Novitsky et al. [38] data, suggesting that a strong enough signal in the data can overcome the prior.

As noted by Lintusaari et al. [31], uniform priors on model parameters may translate to highly informative priors on quantities of interest. We observed a non-linear relationship between the preferential attachment power *α* and the power law exponent *γ* (fig. S9). Therefore, placing a uniform prior on *α* between 0 and 2 is equivalent to placing an informative prior that *γ* is close to 2. Therefore, if we were primarily interested in *γ* rather than *α*, a more sensible choice of prior might have a shape informed by fig. S9 and be bounded above by approximately *α* = 1.5. This would uniformly bound *γ* in the region 2 ≤ *γ* ≤ 4 commonly reported in the network literature [30, 47, 8, 6]. We note however that Jones and Handcock [25] estimated *γ* values greater than four for some datasets, in one case as high as 17, indicating that a wider range of permitted *γ* values may be warranted.

Our investigation of published HIV datasets indicated heterogeneity in the contact network structures underlying several distinct local epidemics. When interpreting these results, we caution that the BA model is quite simple and most likely misspecified for these data. In particular, the average degree of a node in the network is equal to 2*m*, and therefore is constrained to be a multiple of 2. Furthermore, we considered the case *m* = 1, where the network has no cycles, to be implausible and therefore assigned it zero prior probability in one set of analyses. This forced the average degree to be at least four, which may be unrealistically high for sexual networks. The fact that the estimated values of *α* differed substantially for three datasets depending on whether or not *m* = 1 was allowed by the prior is further evidence of this potential misspecification. However, we note that for two of the datasets, the estimated values of *α* did not change much between priors, and the estimates of *I* were robust to the choice of prior for all datasets studied. More sophisticated models, for example models incorporating heterogeneity in node behaviour, are likely to provide a better fit to these data.

With respect to the preferential attachment power *α*, the five datasets analysed fell into three categories (fig. 4). First, we estimated a preferential attachment power close to 1, indicating linear preferential attachment, for the outbreaks studied by Niculescu et al. [36] and Wang et al. [59]. These values were robust to specifying different priors for *m*. Both studies were of populations in which we would expect a high degree of epidemiological relatedness: Niculescu et al. [36] studied a recent outbreak among Romanian injection drug users (IDU), while Wang et al. sampled acutely infected MSM in Beijing, China. Both these are contexts in which we would expect some of the assumptions of the BA model, such as a connected network, relatively high mean degree, and preferential attachment dynamics, to hold.

The remaining three datasets (Cuevas et al. [11], Novitsky et al. [38], and Li et al. [29]) had estimated values of *α* below 0.5 when *m* = 1 was included in the prior, but these were not robust to changing the prior to exclude *m* = 1. For the Cuevas et al. data, model misspecification is likely partially responsible. While the authors found that a large proportion of the samples were epidemiologically linked, these were mainly in small local clusters rather than the single large component postulated by the BA model. In addition, the mixed risk groups in the dataset would be unlikely to significantly interact, further weakening any global preferential attachment dynamics. The dataset studied by Novitsky et al. [38] originated from a densely sampled population where the predominant risk factor was believed to be heterosexual exposure. Although the MAP estimate of *α* was almost unchanged when the value *m* = 1 was excluded from the prior, the confidence interval shrank significantly. For both priors, the estimated prevalence was extremely high, in fact higher than the estimated HIV prevalence in the sampled region. The authors indicated that the source of the samples was a town in close proximity to the country’s capital city, and suggested that there may have been a high degree of migration and partner interchange between the two locations. It is possible that the contact network underlying the subtree we investigated includes a much larger group based in the capital city, which would explain the high estimate of *I*. There is no clear explanation for the discrepancy between the two priors for the Li et al. [29] data, as the subset we analyzed formed a phylogenetic cluster and therefore was a good candidate for the BA model. However, nearly all the posterior density was assigned to *m* = 1 when this value was allowed, indicating that the network was more likely to have an acyclic tree structure.

In addition to the aforementioned possibility of misspecification, additional modelling assumptions include the network being connected and static, all transmission rates being equal, no removal after infection, identical behaviour of all nodes, and random sampling. The last two were addressed with small-scale experiments. We simulated a network where some nodes exhibited a higher attachment power than others, and found that the estimated attachment power was simply the average of the two values. This indicated that, although we could characterize the network in aggregate, the estimated parameters could not be said to apply to any individual node. The effect of biased sampling was investigated by analysing a transmission tree which had been sampled in a peer-driven fashion. The results were roughly in line with those for random sampling, however the estimated value of *α* was lower than the average for randomly-sampled trees. Further experiments would be necessary to fully explore the impact of these assumptions on the method’s accuracy. However, despite these issues, we felt it was best to demonstrate the method first on a simple model. It is possible to use this framework to fit more complex models which address some of these issues, such as one incorporating heterogeneous node behaviour, which may prove a fruitful avenue for future investigations.

Our method has a number of caveats, perhaps the most significant being that it takes a transmission tree as input. In reality, true transmission trees are not available and must be approximated, often by way of a viral phylogeny. Although this has been demonstrated to be a fair approximation [e.g. 27], and is frequently used in practice [e.g. 50], the topologies of a viral phylogeny and transmission tree can differ significantly [61] due to within-host evolution and the sampling process. In addition, the ABC-SMC algorithm is computationally intensive, taking about a day when run on 20 cores in parallel with the settings we described in the methods. Nevertheless, our method is potentially useful to epidemiological researchers interested in the general characteristics of the network structure underlying disease outbreaks. This work, and previous work by our group [41], has demonstrated that kernel-ABC is a broadly applicable and effective framework in which to perform phylodynamic inference.

## Acknowledgements

This work was supported by grants from the Canadian Institutes of Health Research (CIHR, operating grant HOP-111406), and the Bill & Melinda Gates Foundation (award number OPP1110049). R.M.M. was supported by a scholarship from the CIHR Strategic Training Program in Bioinformatics. A.F.Y.P. was supported by a CIHR New Investigator Award (Canadian HIV Vaccine Initiative, Vaccine Discovery and Social Research) and by a Career Investigator Scholar Award from the Michael Smith Foundation for Health Research, in partnership with the Providence Health Care Research Institute and St. Paul’s Hospital Foundation.

## References

[1] Albert-László Barabási and Réka Albert. “Emergence of scaling in random networks”. In Science 286.5439 (1999), pp. 509–512.

[2] Marc Barthélemy et al. “Dynamical patterns of epidemic outbreaks in complex heterogeneous networks”. In Journal of Theoretical Biology 235.2 (2005), pp. 275–288.

[3] Mark A Beaumont, Wenyang Zhang, and David J Balding. “Approximate Bayesian computation in population genetics”. In Genetics 162.4 (2002), pp. 2025–2035.

[4] Mark A Beaumont et al. “Adaptive approximate Bayesian computation”. In Biometrika 96.4 (2009), pp. 983–990.

[5] Tom Britton and Philip D O’Neill. “Bayesian inference for stochastic epidemics in populations with random social structure”. In Scandinavian Journal of Statistics 29.3 (2002), pp. 375–390.

[6] Andrew J Leigh Brown et al. “Transmission network parameters estimated from HIV sequences for a nationwide epidemic”. In The Journal of Infectious Diseases 204.9 (2011), p. 1463.

[7] Peter JA Cock et al. “Biopython: freely available Python tools for computational molecular biology and bioinformatics”. In Bioinformatics 25.11 (2009), pp. 1422–1423.

[8] Stirling A Colgate et al. “Risk behavior-based model of the cubic growth of acquired immunodeficiency syndrome in the United States”. In Proceedings of the National Academy of Sciences 86.12 (1989), pp. 4793–4797.

[9] Caroline Colijn and Jennifer Gardy. “Phylogenetic tree shapes resolve disease transmission patterns”. In Evolution, Medicine, and Public Health 2014.1 (2014), pp. 96–108.

[10] Gabor Csardi and Tamas Nepusz. “The igraph software package for complex network research”. In InterJournal, Complex Systems 1695.5 (2006), pp. 1–9.

[11] MT Cuevas et al. “HIV-1 transmission cluster with T215D revertant mutation among newly diagnosed patients from the Basque Country, Spain”. In Journal of Acquired Immune Deficiency Syndromes 51.1 (2009), p. 99.

[12] Pierre Del Moral, Arnaud Doucet, and Ajay Jasra. “An adaptive sequential Monte Carlo method for approximate Bayesian computation”. In Statistics and Computing 22.5 (2012), pp. 1009–1020.

[13] Alexei J Drummond and Andrew Rambaut. “BEAST: Bayesian evolutionary analysis by sampling trees”. In BMC Evolutionary Biology 7.1 (2007), p. 214.

[14] Alexei J Drummond et al. “Measurably evolving populations”. In Trends in Ecology & Evolution 18.9 (2003), pp. 481–488.

[15] Robert C Edgar. “MUSCLE: multiple sequence alignment with high accuracy and high throughput”. In Nucleic Acids Research 32.5 (2004), pp. 1792–1797.

[16] Paul Erdős and Alfred Rényi. “On the evolution of random graphs”. In Publications of the Mathematical Institute of the Hungarian Academy of Sciences 5 (1960), pp. 17–61.

[17] Daniel T Gillespie. “A general method for numerically simulating the stochastic time evolution of coupled chemical reactions”. In Journal of Computational Physics 22.4 (1976), pp. 403–434.

[18] Steven M Goodreau. “Assessing the effects of human mixing patterns on human immunodeficiency virus-1 interhost phylogenetics through social network simulation”. In Genetics 172.4 (2006), pp. 2033–2045.

[19] Manolo Gouy, Stéphane Guindon, and Olivier Gascuel. “SeaView version 4: a multiplatform graphical user interface for sequence alignment and phylogenetic tree building”. In Molecular Biology and Evolution 27.2 (2010), pp. 221–224.

[20] Bryan T Grenfell et al. “Unifying the epidemiological and evolutionary dynamics of pathogens”. In Science 303.5656 (2004), pp. 327–332.

[21] Chris Groendyke, David Welch, and David R Hunter. “Bayesian inference for contact networks given epidemic data”. In Scandinavian Journal of Statistics 38.3 (2011), pp. 600–616.

[22] Mark S Handcock and James Holland Jones. “Likelihood-based inference for stochastic models of sexual network formation”. In Theoretical Population Biology 65.4 (2004), pp. 413–422.

[23] Gareth J Hughes et al. “Molecular Phylodynamics of the Heterosexual HIV Epidemic in the United Kingdom”. In PLoS Pathogens 5.9 (2009).

[24] Thijs Janzen, Sebastian Höhna, and Rampal S Etienne. “Approximate Bayesian computation of diversification rates from molecular phylogenies: introducing a new efficient summary statistic, the nLTT”. In Methods in Ecology and Evolution 6.5 (2015), pp. 566–575.

[25] James Holland Jones and Mark S Handcock. “An assessment of preferential attachment as a mechanism for human sexual network formation”. In Proceedings of the Royal Society of London B: Biological Sciences 270.1520 (2003), pp. 1123–1128.

[26] Alden S Klovdahl. “Social networks and the spread of infectious diseases: the AIDS example”. In Social Science & Medicine 21.11 (1985), pp. 1203–1216.

[27] Thomas Leitner et al. “Accurate reconstruction of a known HIV-1 transmission history by phylogenetic tree analysis”. In Proceedings of the National Academy of Sciences 93.20 (1996), pp. 10864–10869.

[28] Gabriel E Leventhal et al. “Inferring Epidemic Contact Structure from Phylogenetic Trees”. In PLoS Computational Biology 8.3 (2012).

[29] Xiaoyan Li et al. “HIV-1 Genetic Diversity and Its Impact on Baseline CD4+ T Cells and Viral Loads among Recently Infected Men Who Have Sex with Men in Shanghai, China”. In PLoS ONE 10.6 (2015).

[30] Fredrik Liljeros et al. “The web of human sexual contacts”. In Nature 411.6840 (2001), pp. 907–908.

[31] Jarno Lintusaari et al. “On the identifiability of transmission dynamic models for infectious diseases”. In Genetics (2016).

[32] Susan J Little et al. “Using HIV Networks to Inform Real Time Prevention Interventions”. In PLoS ONE 9.6 (2014).

[33] Junling Ma, P van den Driessche, and Frederick H Willeboordse. “The importance of contact network topology for the success of vaccination strategies”. In Journal of theoretical biology 325 (2013), pp. 12–21.

[34] Martina Morris. “Epidemiology and social networks: modeling structured diffusion”. In Sociological Methods & Research 22.1 (1993), pp. 99–126.

[35] Shigeki Nakagome, Kenji Fukumizu, and Shuhei Mano. “Kernel approximate Bayesian computation in population genetic inferences”. In Statistical Applications in Genetics and Molecular Biology 12.6 (2013), pp. 667–678.

[36] Iulia Niculescu et al. “Recent HIV-1 outbreak among intravenous drug users in Romania: evidence for cocirculation of CRF14_BG and subtype F1 strains”. In AIDS Research and Human Retroviruses 31.5 (2015), pp. 488–495.

[37] Vladimir Novitsky et al. “Phylogenetic relatedness of circulating HIV-1C variants in Mochudi, Botswana”. In PLoS ONE 8.12 (2013).

[38] Vlad Novitsky et al. “Impact of sampling density on the extent of HIV clustering”. In AIDS Research and Human Retroviruses 30.12 (2014), pp. 1226–1235.

[39] Eamon B O’Dea and Claus O Wilke. “Contact heterogeneity and phylodynamics: how contact networks shape parasite evolutionary trees”. In Interdisciplinary Perspectives on Infectious Diseases (2011).

[40] Martyn Plummer et al. “CODA: Convergence diagnosis and output analysis for MCMC”. In R News 6.1 (2006), pp. 7–11.

[41] Art FY Poon. “Phylodynamic inference with kernel ABC and its application to HIV epidemiology”. In Molecular Biology and Evolution 32.9 (2015), pp. 2483–2495.

[42] Art FY Poon et al. “Mapping the shapes of phylogenetic trees from human and zoonotic RNA viruses”. In PLoS ONE 8.11 (2013).

[43] Art FY Poon et al. “The impact of clinical, demographic and risk factors on rates of HIV transmission: a population-based phylogenetic analysis in British Columbia, Canada”. In The Journal of Infectious Diseases 211.6 (2015), pp. 926–935.

[44] Morgan N Price, Paramvir S Dehal, and Adam P Arkin. “FastTree 2–approximately maximum-likelihood trees for large alignments”. In PloS one 5.3 (2010), e9490.

[45] David A Rasmussen, Erik M Volz, and Katia Koelle. “Phylodynamic inference for structured epidemiological models”. In PLoS Computational Biology 10.4 (2014).

[46] Katy Robinson et al. “How the dynamics and structure of sexual contact networks shape pathogen phylogenies”. In PLoS Computational Biology 9.6 (2013).

[47] Anne Schneeberger et al. “Scale-free networks and sexually transmitted diseases: a description of observed patterns of sexual contacts in Britain and Zimbabwe”. In Sexually Transmitted Diseases 31.6 (2004), pp. 380–387.

[48] Kwang-Tsao Shao. “Tree balance”. In Systematic Biology 39.3 (1990), pp. 266–276.

[49] Scott A Sisson, Yanan Fan, and Mark M Tanaka. “Sequential Monte Carlo without likelihoods”. In Proceedings of the National Academy of Sciences 104.6 (2007), pp. 1760–1765.

[50] Tanja Stadler and Sebastian Bonhoeffer. “Uncovering epidemiological dynamics in heterogeneous host populations using phylogenetic methods”. In Philosophical Transactions of the Royal Society B: Biological Sciences 368.1614 (2013).

[51] Tanja Stadler et al. “Estimating the basic reproductive number from viral sequence data”. In Molecular Biology and Evolution 29.1 (2012), pp. 347–357.

[52] Mikael Sunnåker et al. “Approximate Bayesian Computation”. In PLoS Computational Biology 9.1 (2013).

[53] Simon Tavaré et al. “Inferring coalescence times from DNA sequence data”. In Genetics 145.2 (1997), pp. 505–518.

[54] Luc Villandre et al. “Assessment of overlap of phylogenetic transmission clusters and communities in simple sexual contact networks: applications to HIV-1”. In PloS ONE 11.2 (2016).

[55] Erik Volz. “SIR dynamics in random networks with heterogeneous connectivity”. In Journal of Mathematical Biology 56.3 (2008), pp. 293–310.

[56] Erik M Volz. “Complex population dynamics and the coalescent under neutrality”. In Genetics 190.1 (2012), pp. 187–201.

[57] Erik M Volz et al. “Simple epidemiological dynamics explain phylogenetic clustering of HIV from patients with recent infection”. In PLoS Computational Biology 8.6 (2012).

[58] Erik Volz and Lauren Ancel Meyers. “Susceptible–infected–recovered epidemics in dynamic contact networks”. In Proceedings of the Royal Society of London B: Biological Sciences 274.1628 (2007), pp. 2925–2934.

[59] X Wang et al. “Targeting HIV prevention based on molecular epidemiology among deeply sampled subnetworks of men who have sex with men”. In Clinical Infectious Diseases 61.9 (2015), p. 1462.

[60] David Welch, Shweta Bansal, and David R Hunter. “Statistical inference to advance network models in epidemiology”. In Epidemics 3.1 (2011), pp. 38–45.

[61] Rolf JF Ypma, W Marijn van Ballegooijen, and Jacco Wallinga. “Relating phylogenetic trees to transmission trees of infectious disease outbreaks”. In Genetics 195.3 (2013), pp. 1055–1062.

[62] Achim Zeileis et al. “kernlab-an S4 package for kernel methods in R”. In Journal of Statistical Software 11.9 (2004), pp. 1–20.

